# Detection of differential RNA modifications from direct RNA sequencing of human cell lines

**DOI:** 10.1101/2020.06.18.160010

**Authors:** Ploy N. Pratanwanich, Fei Yao, Ying Chen, Casslynn W.Q. Koh, Christopher Hendra, Polly Poon, Yeek Teck Goh, Phoebe M. L. Yap, Choi Jing Yuan, Wee Joo Chng, Sarah Ng, Alexandre Thiery, W.S. Sho Goh, Jonathan Göke

## Abstract

Differences in RNA expression can provide insights into the molecular identity of a cell, pathways involved in human diseases, and variation in RNA levels across patients associated with clinical phenotypes. RNA modifications such as m6A have been found to contribute to molecular functions of RNAs. However, quantification of differences in RNA modifications has been challenging. Here we develop a computational method (xPore) to identify differential RNA modifications from direct RNA sequencing data. We evaluate our method on transcriptome-wide m6A profiling data, demonstrating that xPore identifies positions of m6A sites at single base resolution, estimates the fraction of modified RNAs in the cell, and quantifies the differential modification rate across conditions. We apply the method to direct RNA-Sequencing data from 6 cell lines and find that many m6A sites are preserved, while a subset of m6A sites show significant differences in their modification rates across cell types. Together, we show that RNA modifications can be identified from direct RNA-sequencing with high accuracy, enabling the analysis of differential modifications and expression from a single high throughput experiment.

**Availability:** xPore is available as open source software (https://github.com/GoekeLab/xpore)

## Introduction

Each cell, tissue, and organ is defined through the set of genes that are expressed. Yet, it’s not just the level of transcription that defines the molecular profile of the cell: more than 100 modifications of RNAs have the potential to alter cellular mechanisms and modulate disease risks. Among the known RNA modifications, the most abundant mRNA modification is N6-methyladenosine (m6A), with an average of one to three m6A modified sites per transcript (Dominissini et al. 2012; Meyer et al. 2012), that impacts several post-transcriptional processes including mRNA decay, mRNA translation, pre-mRNA splicing, and pri-miRNA processing (Zaccara, Ries, and Jaffrey 2019). RNA modifications have been found to be essential during early development (Zheng et al. 2013; Wang et al. 2014; Zhao et al. 2014; Geula et al. 2015; Chen et al. 2015; Xu et al. 2017) and aberrant modifications have been associated with diseases (Mathiyalagan et al. 2019; Z. Li et al. 2017; Su et al. 2018; Deng et al. 2018). In cancer, the first RNA therapeutics that target RNA modifying enzymes are being developed, highlighting their impact on precision medicine. A comprehensive profiling of the transcriptome therefore involves the quantification of both transcript levels and modification rates.

Quantification of transcript expression is done by sequencing cDNA (“RNA-Seq”), one of the most widely used assays in molecular biology. In contrast, identifying and quantifying RNA modifications still remains a major challenge. The most common modifications (m6A, ac4C, m5C and hm5C) can be mapped using short-read cDNA sequencing-based methods (Zaccara, Ries, and Jaffrey 2019). For m6A, cross-linking-immunoprecipitation (CLIP)-based methods map these modifications transcriptome-wide (Linder et al. 2015), and DART-Seq (Meyer 2019) and m6ACE-Seq (Koh, Goh, and Sho Goh 2019) additionally enable quantifying the modification rate of individual m6A sites. However, cDNA based methods use reverse transcription which might introduce biases; they rely on available antibodies or known enzymes that limit the profiling of most modifications, and the requirement for specialised protocols prevents their large-scale application.

Third generation sequencing using the Oxford Nanopore technology promises to overcome these limitations through direct sequencing of native RNA (direct RNA-seq/ dRNA-Seq) (Garalde et al. 2018). Direct RNA-Seq applies a fundamentally unique principle for base identification: as RNA passes through the pore, the magnitudes of electric intensity across the nanopore surface are recorded and used to identify the corresponding nucleotide sequence. RNA modifications cause shifts in the intensity levels which are used to computationally identify modified bases (H. Liu et al. 2019; Lorenz et al. 2020; Leger et al. 2019; Stoiber et al. 2017). Identification of m6A modifications can be achieved using a supervised learning approach that heavily relies on basecalling accuracy or training data from synthetic sequences (H. Liu et al. 2019). An alternative approach is the detection of modifications by comparison to a non-modified control sample, thereby removing the requirements of training data, and potentially enabling the identification of non-m6A modifications (Stoiber et al. 2017; Leger et al. 2019). However, such samples are difficult to generate, they are affected by artifacts from various depletion methods, require prior knowledge about enzymes, and are still influenced by non-depleted modifications.

One of the most frequent applications for transcriptome profiling is the analysis of differential expression. Here, we present an analogous statistical framework that defines measures of significance and effect size for differential RNA modifications from direct RNA sequencing data, as such removing the strict requirement for a non-modified training sample and enabling the simultaneous profiling of differential transcript expression and modification without any additional experiments. To determine the proportion of RNA modifications across multiple samples, we fit a multi-sample two-Gaussian mixture model, and infer directionality of modification rate differences by utilizing information across all tested positions. We evaluate our method transcriptome-wide in a loss-of-m6A system after knockout of *Mettl3*. We then applied our method on direct RNA-Seq data from 6 of the most commonly used human cell lines including cancer tissues, providing insights into the dynamics of m6A. Our study introduces a computational method that enables the profiling of differential RNA modifications transcriptome wide and provides the largest currently available direct RNA-Seq data, which will be valuable as a resource and benchmark data set.

## Results

### xPore: Detection of differential RNA modifications from direct RNA-Seq data

Nanopore Direct RNA-Sequencing generates a raw ionic current signal for each individual read (Figure 1A). During the sequencing process, the nanopores measure a signal corresponding to an RNA sequence of length 5 that resides in the pore, and a signal shift is observed when the next base enters the pore (Figure 1B). Here we use the normalised mean signal information at each 5-mer (*event*) obtained after transcriptome alignment with Minimap2 (H. Li 2018) and signal segmentation using Nanopolish Eventalign (Loman, Quick, and Simpson 2015; Simpson et al. 2017).

**Figure 1.**
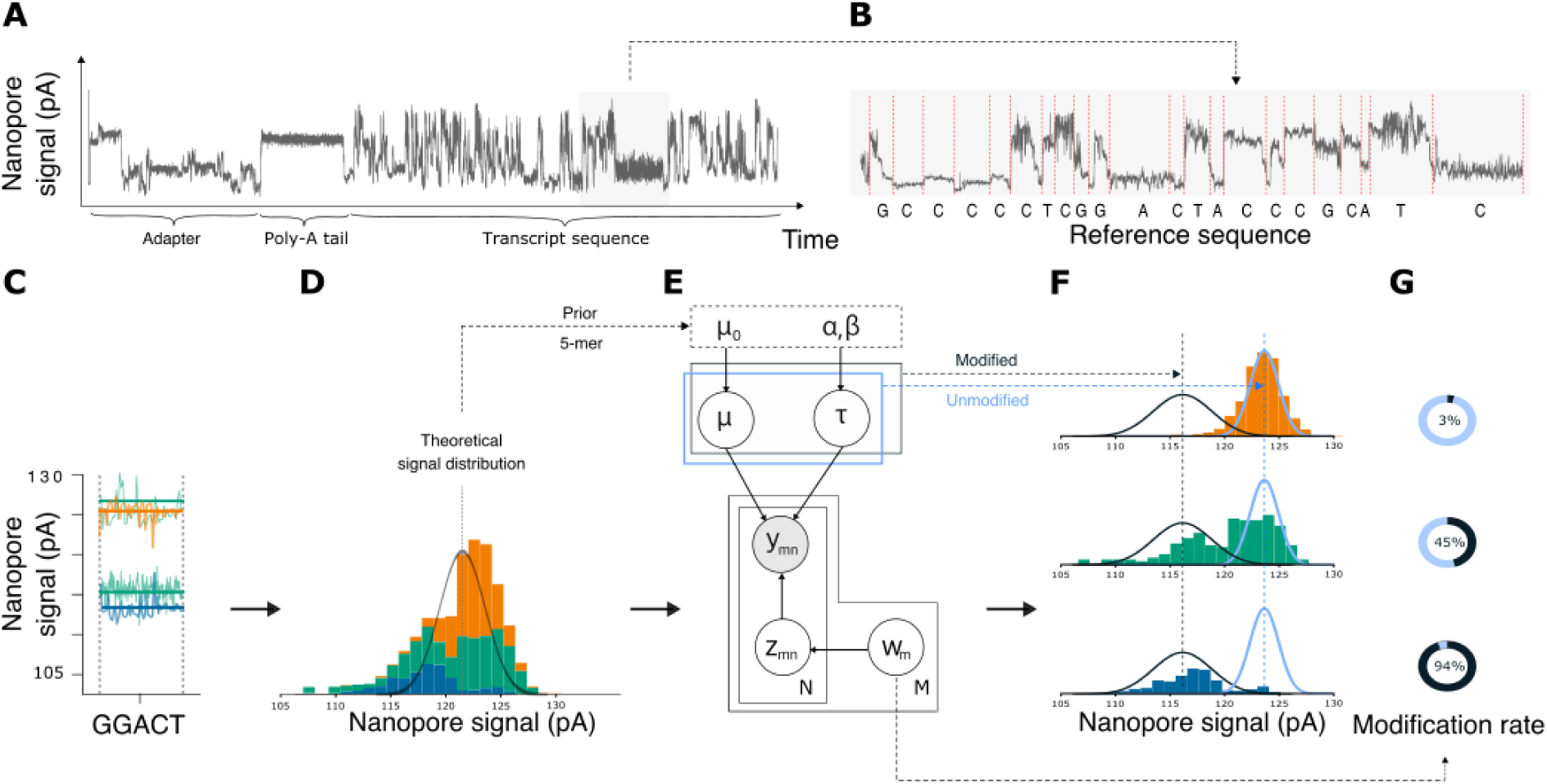
Quantification of RNA modifications from direct RNA-Seq data using xPore. (A) Example of raw signal data from a direct RNA sequencing read. (B) A close-up look of the raw signal with the corresponding transcript sequence obtained from basecalling, sequence alignment, and signal segmentation. (C) Signal of multiple reads aligned at a GGACT site from different samples (orange, green, and blue) (D) Shown is a histogram of the mean signal from all reads covering a position for three different samples (orange, green, and blue). The black line indicates the expected distribution for unmodified RNA, samples which contain modified RNAs will show a bimodal distribution. (E) Graphical representation of the model used by xPore to quantify the modification rate at each position. The gray circle indicates observed variables (data), the white circles indicate unobserved variables that are estimated by xPore. (F) xPore estimates the parameters for two Gaussian distributions corresponding to modified (black) and unmodified RNAs (blue). (G) xPore summarises the modification rates for each sample.

In order to detect differential RNA modification rate for each genomic position across samples (Figure 1C), we developed xPore, a computational method that analyses dRNA-Seq data at the signal level. xPore models a mixture of two Gaussian distributions, corresponding to unmodified and modified RNAs. To guide the parameter estimation, we impose the theoretical signal distribution of unmodified RNA as a prior on the two Gaussians (Figure 1D; see Methods). The means and variances of these two distributions are modelled to be shared across samples (Figure 1E; see Methods). After a few iterations of variational Bayesian inference (Corduneanu and M. 2001), inferred means and variances are obtained (Figure 1F). We then assign the inferred distribution that is closer to the theoretical mean as unmodified, assigning the other distribution as modified. xPore also learns the modification probability of each read, which allows us to compute the fraction of modified reads as an estimate of the modification rate per sample (Figure 1G).

To increase the precision and control the number of false positives, we implemented two filtering steps. Firstly, we exclude positions where the distributions for the unmodified and modified signals are nearly identical, thereby reducing the number of tests and increasing the power. Secondly, RNA modifications induce a systematic shift in the signal for each k-mer, as the same RNA modification will either increase or decrease the signal, but not both. Although xPore does not identify the type of modification at each position, we can nevertheless restrict the analysis to a single modification at each k-mer by only considering one-directional signal shifts. This filter removes outliers and enables the transcriptome-wide comparison of modification patterns. On the remaining sites, we perform a statistical test on the differential modification rates between samples and prioritise differentially modified sites accordingly.

The method is implemented in Python and available as part of the open source package xPore on github (https://github.com/GoekeLab/xpore). Using FAST5 files as input, xPore returns a table summarizing the means and variances of the unmodified and modified distributions, the assignment confidence levels, the modification rates for each sample, and the test statistics for individual positions.

### xPore accurately identifies m6A sites at single base resolution

One of the most abundant and best studied RNA modifications is m6A (Zaccara, Ries, and Jaffrey 2019). To evaluate the ability of our method to detect differentially modified sites in the human transcriptome, we compared wild type HEK293T cells (‘wildtype cells’) with cells where expression of the main m6A writer, Mettl3 was deleted via CRISPR-Cas9 (‘knockout cells’). We generated 3 replicates for both the wildtype and knockout cells, resulting in nearly 8 million reads in total. After filtering out low coverage positions, we had over 9 million sites to be modelled, 939,902 of which were passed through the post-modelling filter and were tested for differential modifications. For evaluation, we used single-base-resolution m6ACE-Seq and the presence of the DRACH motif to estimate the number of correctly predicted sites (Figure 2A).

**Figure 2.**
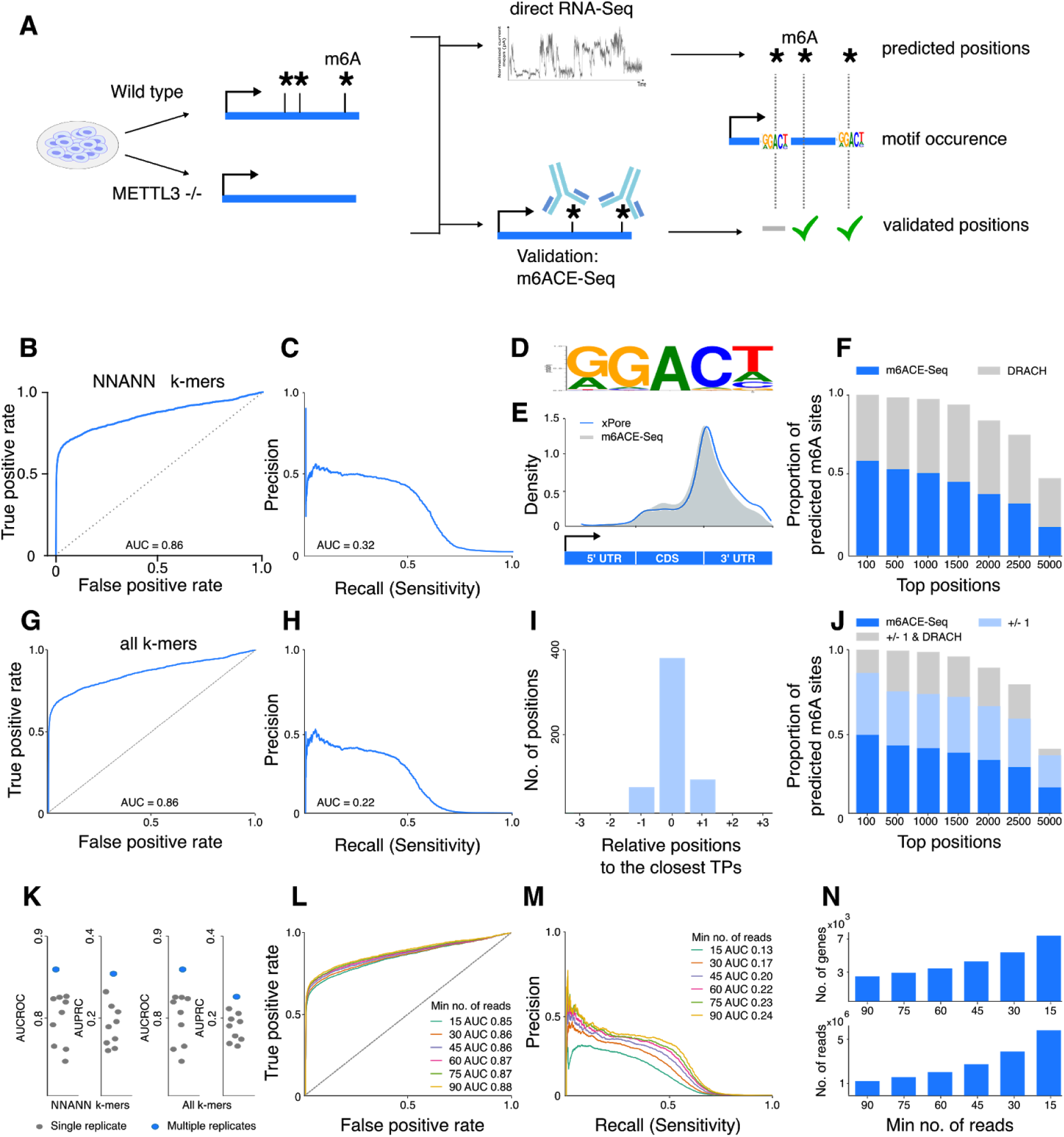
Detection of m6A sites in the human transcriptome. (A) Experimental design: RNA of wildtype and *Mettl3*-knockout HEK293T cells is sequenced by direct RNA-Seq and differential m6A modification rates between both cells are estimated using xPore. The results are validated against the results from m6ACE-Seq and tested for the occurence of the m6A motif (DRACH). (B) ROC curve and (C) Precision Recall curve for candidate m6A sites identified by xPore using the set of m6ACE-Seq sites as ground truth. Only kmers with an A at the center are used in this analysis. (D) K-mers from candidate m6A sites identified by xPore resemble the expected m6A motif. (E) Candidate m6A sites identified by xPore are enriched at the 3’ end of the CDS, resembling the expected distribution shown for m6ACE-Seq results. (F) Proportion of predicted m6A sites that overlap with m6ACE-Seq sites (dark blue) and which resemble the DRACH motif (gray). (G) ROC curve and (H) Precision Recall curve for candidate m6A sites identified by xPore using the set of m6ACE-Seq sites as ground truth. All kmers are used in this analysis. (I) Relative position of predicted m6A sites to the closest m6ACE-Seq validated site (true positives (TPs)). (J) Proportion of predicted m6A sites that overlap with m6ACE-Seq sites (dark blue), which are within 1 base distance to a m6ACE-Seq site (light blue), and which resemble the DRACH motif (gray). (K) Area under ROC and precision-recall curves when 3 replicates are used (blue) compared to a single replicate analysis (gray). Left: kmers with A in the center. RIght: all kmers. (L) ROC curve and (M) Precision Recall curve when the direct RNA-Seq data from multiple replicates are pooled. Using different thresholds for the minimum number of reads influences the sensitivity and precision. (N) Read coverage of genes (bottom) and number of genes that can be analysed at these thresholds. xPore can analyse more than 7,000 genes at a minimum read coverage of 15.

Firstly, we evaluated the ability to identify differentially m6A modified sites at positions with the base A (NNANN). Among all tested positions, xPore achieves an overall AUC of 0.86 when m6ACE-Seq is used as a reference (Figure 2B). As the number of unmodified positions is much larger than the number of modified positions, we particularly investigated the precision of our predictions at different levels of sensitivity (Figure 2C). The model has a precision of 0.60 at the top predictions when m6ACE-Seq is used as m6A reference. Strikingly, many of the top ranked positions by xPore that are not identified as modified by m6ACE-Seq still showed the DRACH motif (Figure 2D), suggesting that Nanopore direct RNA-Seq might help in identifying a different set of modified sites that had been otherwise missed by antibody-based detection methods. One of the hallmarks of m6A sites is the distribution along the transcript that is specifically enriched close to the stop codon (Koh, Goh, and Sho Goh 2019; Linder et al. 2015). The top ranked positions are clustered as expected of m6A sites, further suggesting that the top m6A positions identified by xPore contain only a small number of false positives (Figure 2E). Indeed, when we combined m6ACE-Seq data and motif occurrences, xPore achieves an accuracy of >90% among the top significant ∼1,500 positions (p < 0.001), indicating that our method can successfully identify differential m6A modifications even in a search space that covers hundreds of thousands of positions (Figure 2F).

Next we tested the accuracy to identify differentially modified sites when the sequence context is unknown. For this evaluation we ranked transcriptome-wide differentially modified sites in the knockout and wild type cells, including A and non-A nucleotide positions (all k-mers), and again evaluated the performance using m6ACE-Seq and motif content. Overall, xPore achieves an AUC of 0.86 (Figure 2G). Compared to the A-nucleotide analysis, we observed a lower precision at the same level of sensitivity (Figure 2H). A number of m6A sites which are labeled as false positives were neighbouring positions next to modified m6A sites, likely caused by a signal shift due to the modification (Figure 2I). Among the top 1,500 positions, 90% were within a single base distance of a DRACH motif or a validated m6A site, demonstrating that an unbiased search for differential modifications still provides results with high precision among the top ranking positions (Figure 2J).

### Replicates increase precision

Here we analysed data using biological replicates, which might not always be available. To evaluate the performance of xPore in the absence of replicates, we tested every pair of HEK293T wildtype and knockout cells, resulting in 9 pairwise comparisons. Single replicate results were generally less accurate and showed higher variation compared to the multi-replicate analysis (Figure 2K). The results suggest that even in the absence of replicates xPore can prioritise differentially modified sites, however, replicates, which naturally account for biological variation, are recommended to obtain more precise results.

### Pooling data increases sensitivity

The number of positions that can be tested is limited by the sequencing depth of each sample, with lowly expressed genes being potentially excluded. To maximise the number of genes that are modeled by xPore, we therefore combined reads across replicates within the same condition. Using the pooled data we evaluated the performance for different read coverage thresholds. As expected, we observe that a lower read coverage threshold reduces the precision (Figure 2L, M). However, a threshold of 30 reads per position increased the number of genes being tested to more than 5,000, and a threshold of 15 enables the analysis of more than 7,000 genes (Figure 2N). While the analysis using individual replicates with a higher threshold has higher precision, pooling data increases the number of genes which are tested for differential modifications, enabling the detection of RNA modifications at genes which are even lowly expressed.

### Quantitative estimation of cellular RNA modification levels from direct RNA-Seq data

The proportion of RNA molecules which are modified (the *stoichiometry*, or *modification rate*) can show high levels of variation across positions and samples. While some sites are modified in all RNAs, others seem to affect only a minor fraction (N. Liu et al. 2013). However, quantitatively estimating these modification rates remains one of the major challenges (Zaccara, Ries, and Jaffrey 2019). xPore was designed to intrinsically model the probability for each read of being modified. This property enables us to calculate the fraction of reads that are assigned to the modified signal distribution, directly providing an estimate of cellular RNA modification rates.

To evaluate whether the estimated modification rates from xPore represent the true proportion of modified reads, we generated additional RNA mixtures with known expected modification rates. In order to achieve this, we combined various proportions of wildtype and *Mettl3* knockout cellular RNA before profiling them using direct RNA-Seq (Figure 3A, B). Since not all positions are modified at 100% in wild type cells, we investigated the expected relative modification rates, using wild type cells and knockout cells as the reference points. On average, the estimated modification rates from xPore were within 7% of the expected modification rate from the RNA mixtures (Figure 3C). Yet even for individual sites, xPore accurately identifies differentially modified positions and modification rates across replicates (Figure 3D-G). These results indicate that xPore is able to quantitatively estimate the proportion of modified RNAs from direct RNA-Seq data at single-base resolution.

**Figure 3.**
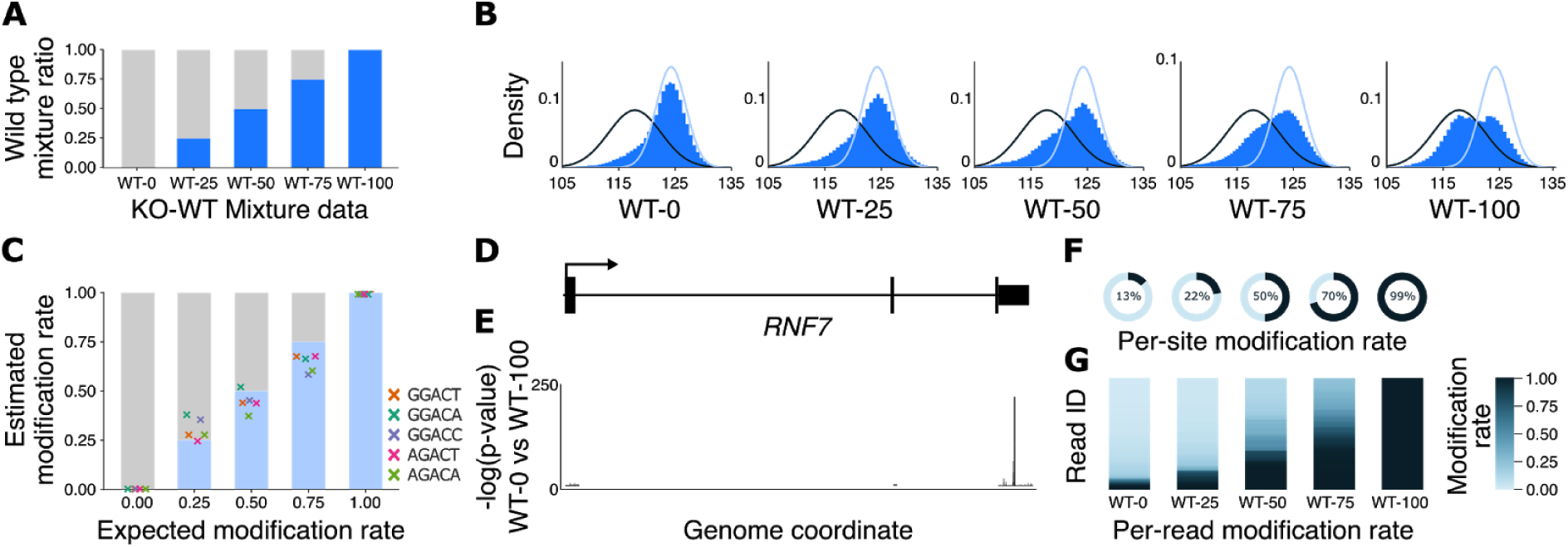
xPore modification rate estimates correspond to the fraction of modified RNAs in the cell. (A) Percentage of wildtype and *Mettl3*-knockout HEK293T cells in each mixture sample. (B) Current intensity level means across all GGACT positions. The black line shows the estimated signal distribution for modified reads from xPore, the blue line shows the estimated signal distribution for unmodified reads. (C) Estimated relative modification rates of the most frequently modified m6A motifs across all modified positions identified by xPore, shown for the different mixture samples. (D) Example of a protein coding gene, *RNF7*, and (E) the −log(p-value) obtained from the pairwise comparison of WT-0 and WT100 showing the most significant differentially m6A site. (F) Per-site modification rate estimates of each mixture sample. (G) Per-read modification rates at the identified GGm6ACT site of *RNF7*.

### Differential modification rates provide an interpretable estimate of effect size

The ability to estimate modification rates allows us to not just identify positions that are modified, but also to quantify *differential* modification rates across conditions. Here we define the *differential modification rate (DMR)* as the difference between the modification rates observed within each condition. Using the expected signal of unmodified RNAs as a reference, we then infer the directionality of the change, allowing us to discriminate positions that show gain and loss of RNA modifications across conditions. As the modification rates correspond to the fraction of modified RNAs in the cell, the differential modification rate can directly be interpreted as a quantitative estimate of effect size. This property enables us to study the effect of any experimental design on the stoichiometry of RNA modifications.

To demonstrate how the DMR can be interpreted, we firstly compared the HEK293T knockout and wild type cells. We identify 1,923 significantly differentially modified positions at NNANN (p<0.001, DMR>0.5) (Figure 4A). Among these, more than 90% are m6A DRACH motifs (Figure 4B). A transcriptome-wide comparison of the modification rates for the most frequently changed k-mers demonstrates the ability to quantitatively identify RNA modifications from direct RNA-Seq data: modification rates within replicates are highly similar (correlation coefficient >0.9 for wild type cells), whereas a systematic increase in modified sited can be observed across conditions (Figure 4C).

**Figure 4.**
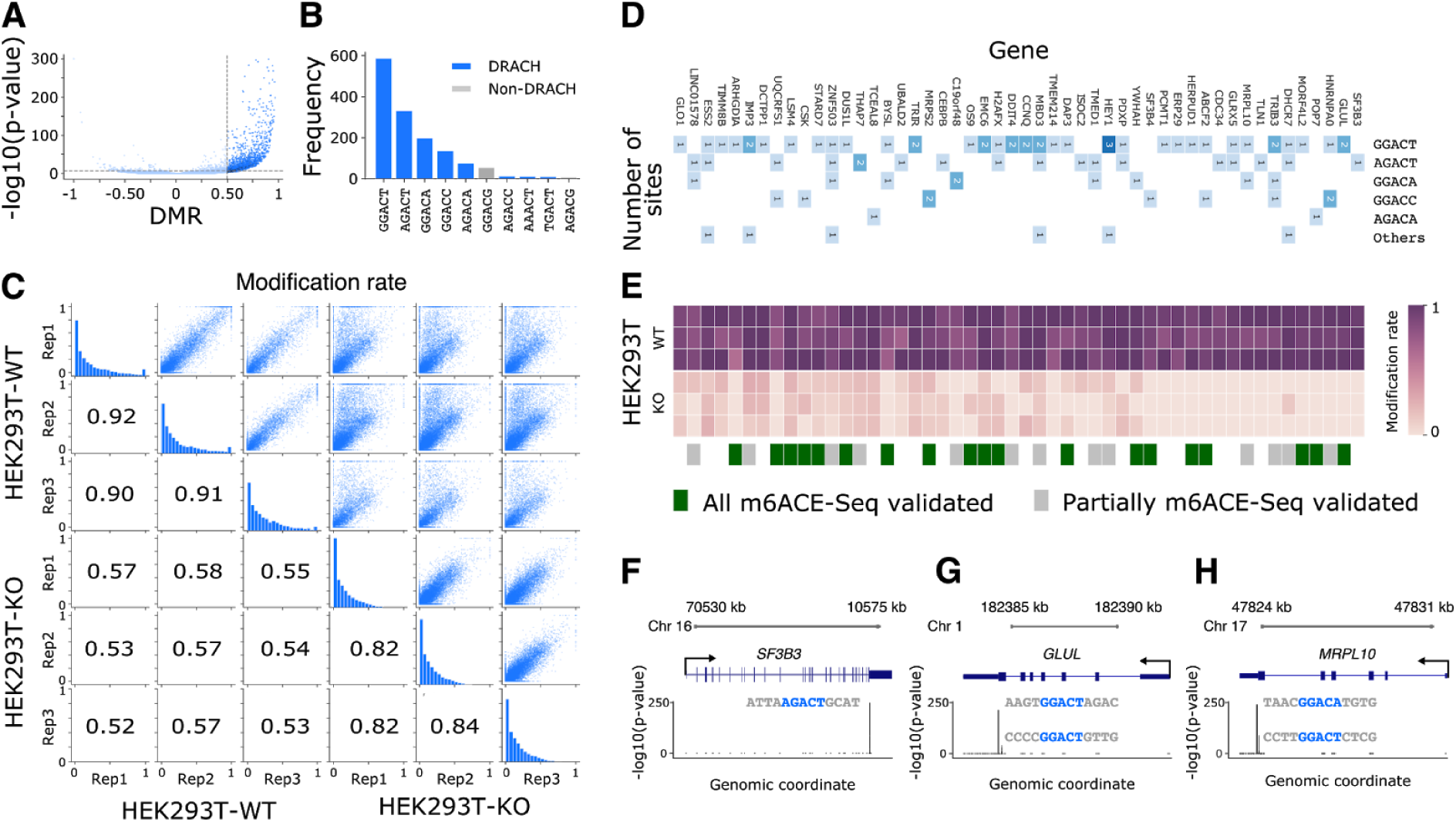
Transcriptome-wide identification of differentially modified positions. (A) Shown is the p-value and the differential modification rate (DMR) for the comparison of HEK293T-WT and HEK293T-KO cells at A-centered k-mers. (B) Frequency of the top 10 k-mers at significantly differentially modified positions. (C) Scatterplot comparing the modification rate estimates for the HEK293T WT and KO samples, histogram of the distribution of modification rates, and pairwise correlation coefficients. (D) The number of modified A sites of the top significant genes ranked by the DMR, along with (E) the corresponding modification rates estimated by xPore across the HEK23T samples. The identified differentially modified sites were all confirmed by m6ACE-Seq in some genes (green), and partially confirmed with newly identified A-modified sites in others (gray). (F-H) Examples of the top ranked genes with the corresponding p-values and the transcript sequence for the identified m6A sites.

Next we identified the set of genes which show the strongest change in RNA modifications after knockout of *Mettl3*. We ranked genes by the highest DMR that was consistently found across replicates (Figure 4D). Many genes appear to contain multiple m6A modifications, frequently involving different kmers (Figure 4D). Many of these positions were confirmed by m6ACE-seq, however, xPore identifies a number of novel positions (Figure 4D, E). Among the genes that are heavily modified by m6A are genes related to RNA processing such as *SF3B3* (Figure 4F), the glutamine synthetase family such as *GLUL* (Figure 4G), and ribosomal proteins such as *MRPL10* (Figure 4H).

### Direct RNA-Sequencing identifies m6A across genetically diverse cell lines

RNA modifications can be detected from direct RNA-Seq as they induce a systematic shift in the signal. However, genetic variants will similarly induce a change in the signal, possibly confounding the results. This effect can be avoided by comparing cell lines which are genetically similar as is often the case in perturbation experiments. To test if xPore can identify differential RNA modifications in samples with a genetically different background, we compared the HEK293T-KO cells against 5 wild type cell lines form the SG-NEx project (https://github.com/GoekeLab/sg-nex-data) covering liver cancer cells (HEPG2), colon cancer cells (HCT116), breast cancer cells (MCF7), lung adenocarcinoma cells (A549), and Leukemia cells (K562) (Figure 5A). To identify RNA modification changes, we focused on the set of RNAs that were expressed across conditions. When we looked into the modification rates at the DRACH motif, loss of *Mettl3* dominated the comparison (Figure 5B). For all cell lines, we can identify between 800 and 2,000 differentially modified sites compared to the HEK293T-KO cells, the vast majority (71% on average) are the m6A DRACH motif (Figure 5C). These results demonstrate that RNA modifications can be identified across conditions even when samples have a distinct genetic background.

**Figure 5.**
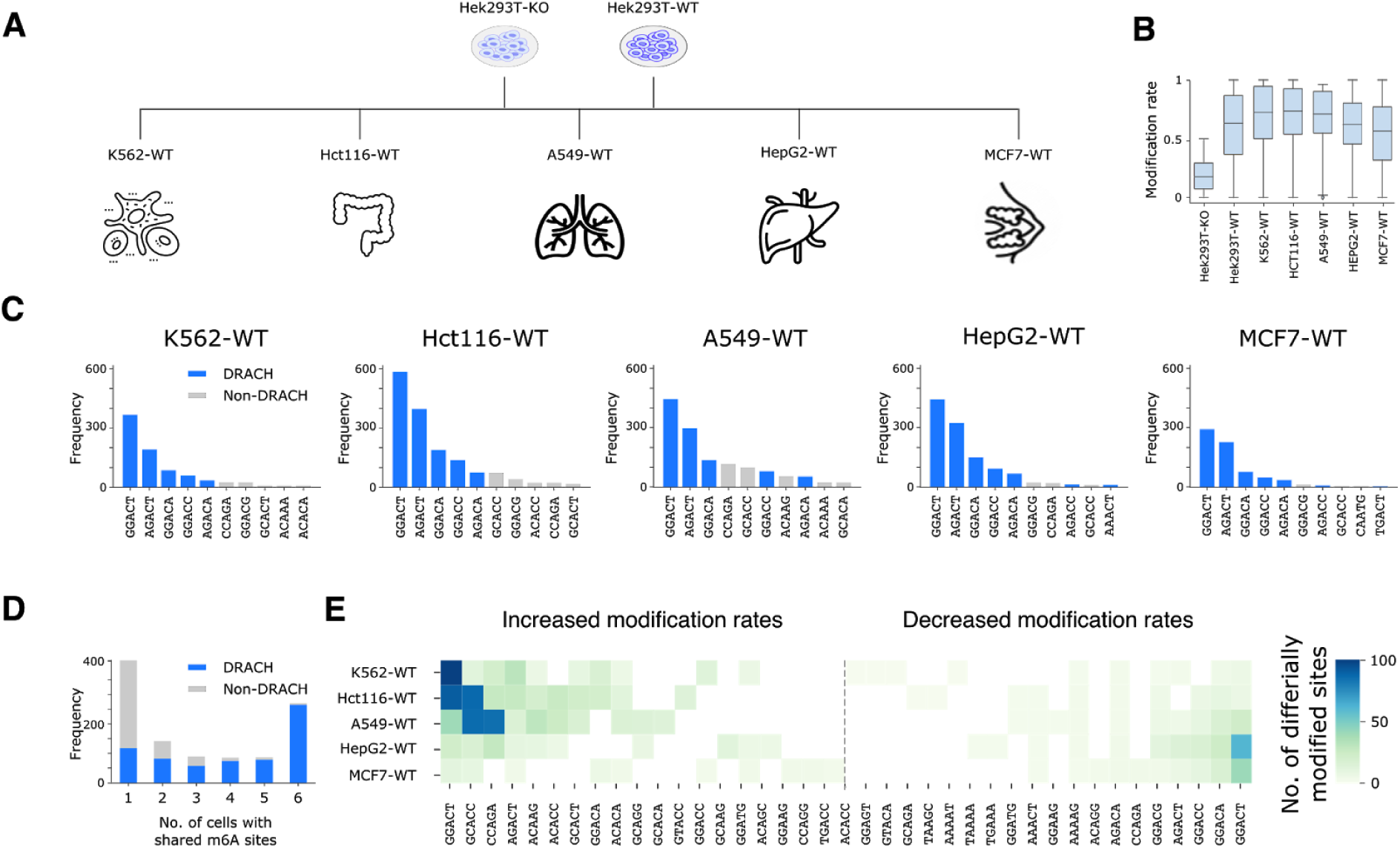
Identification of m6A sites across different tissues and cell lines. (A) Experimental design illustrating the pairwise comparison of each cancer cell line against HEK293T-KO and HEK293T-WT to detect m6A sites and estimate the DMRs. (B) Frequency of the k-mers at the significantly differentially modified sites for each cancer cell line compared to HEK293T-KO cells. K-mers which are classified into DRACH motifs are shown in blue and non-DRACH motifs in gray. (C) Number of m6A sites identified in a single sample and over different number of samples. Most DRACH m6A sites are shared across multiple samples. (D) Average modification rates of m6A identified sites. (E) Heatmap showing the number of significantly differentially modified k-mers for each cancer cell line compared to HEK293T-WT cells: increased modification rate (left) and decreased modification rates (right).

### Variation of m6A across different tissue

It has been shown that m6A modifications differ across tissues and developmental stages, yet quantifying such changes has been challenging. Using the direct RNA-Seq data from the SG-NEx project, we investigated the dynamics of m6A across the different tissues represented by the cell lines. Globally, we find that m6A is stable across cell lines with most positions being shared (Figure 5D). While complete loss and gain of m6A is rare, we observe that a number of m6A sites show quantitative differences between cells. Interestingly, the global modification rate for m6A appears to differ between cells, with K562 showing the highest number of modified m6A sites compared to the other cell lines (Figure 5E). Together, these data indicate that m6A is often stable across tissues, that variation in m6A can be seen at a subset of sites, and that such differences tend to be often quantitative, highlighting the importance to estimate modification rates when comparing RNA modifications across cells or conditions

### Identification of m6A in clinical cancer samples

Clinical samples, in addition to having higher genetic diversity, are often limited by the amount of RNA that can be extracted. For high quality direct RNA-Sequencing, 500ng of polyA RNA is recommended, which can require more than 50mg of total RNA. To test if RNA modifications can be identified in genetically diverse clinical samples with a low amount of RNA, we generated direct RNA-Seq data from 3 multiple myeloma patient samples using only 5% of the recommended RNA amount (2.5ug) (Figure 6A). In total, we obtained more than 1.8 million reads. When we compared these clinical samples to the *Mettl3*-KO cell line data, we observed that a lower number of positions are identified as significantly different relative to cell line samples. However, the top positions are similarly enriched in the DRACH motif (Figure 6B, C), enabling the analysis of m6A modified genes in the multiple myeloma samples (Figure 6D, E). These data suggest that direct RNA-Seq data can be used to identify differential RNA modifications even with genetic diverse conditions and limited RNA, opening opportunities to analyse clinical samples on a larger scale.

**Figure 6.**
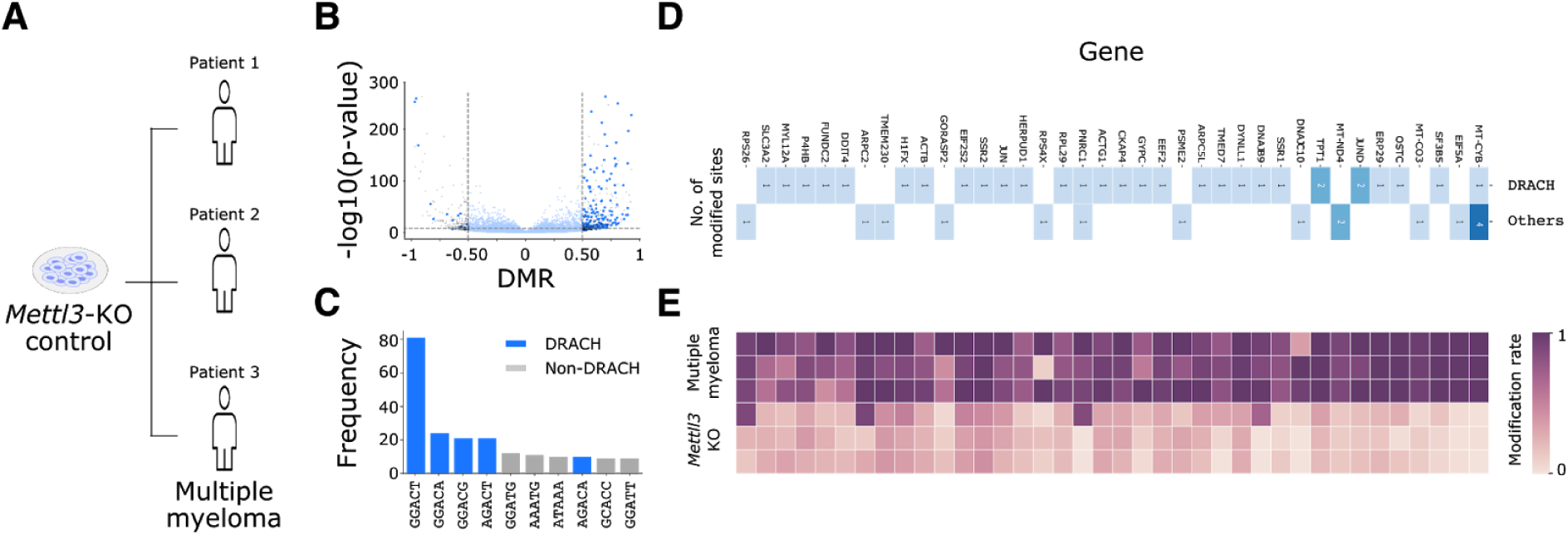
Identification of m6A in clinical samples using direct RNA-Seq. (A) RNA from the *Mettl3*-KO control from HEK293T cells samples and the clinical myeloma samples from three patients are modelled using xPore. (B) Shown is the p-value and the differential modification rate (DMR) for the comparison of *Mettl3*-KO and multiple myeloma samples at A-centered k-mers. (C) Frequency of the top 10 k-mers at significantly differentially modified positions. (D) Number of modified A sites of the top significant genes ranked by the DMR, along with (E) the corresponding modification rates estimated by xPore across the *Mettl3*-KO and multiple myeloma samples.

## Discussion

The transcriptome is one of the major factors that define cellular identity. The differential analysis of transcription is frequently used to understand cellular states, the impact of perturbation experiments, and alterations due to diseases. Here we introduce xPore, a computational method that enables the analysis of differential RNA modifications from direct RNA-Sequencing data, opening a new layer of information that complements transcript expression profiles.

Experimental methods to detect RNA modifications such as m6A have been developed using different approaches (Dominissini et al. 2012; Meyer et al. 2012; Linder et al. 2015; Ke et al. 2015; Garcia-Campos et al. 2019; Meyer 2019; Koh, Goh, and Sho Goh 2019; Shu et al. 2020). Some of the most recent methods enable the identification of thousands of sites at base-resolution, and allow the quantification of modification rates (Garcia-Campos et al. 2019). The main limitation of these approaches is the requirement of sometimes extensive experimental procedures. In contrast, direct RNA-Sequencing promises to enable the analysis of RNA modifications from a single sequencing experiment (Garalde et al. 2018). Methods using direct RNA-Seq have demonstrated the ability to identify m6A and other modifications. However, they often have specific requirements such as a control sample, or do not quantify the stoichiometry of RNA modifications (Stoiber et al. 2017; Leger et al. 2019). xPore achieves base resolution detection of m6A while estimating the modification rate, enabling the quantitative comparison of samples across conditions.

The positions that are identified by xPore show a strong enrichment in the m6A DRACH motif and high validation rates with independent protocols, indicating high precision of predicted positions. Globally, xPore predicts a smaller number of m6A sites compared to other experimental approaches, partially due to stringent filtering that avoids false positives. With increased sequencing throughput and additional replicates, the number of predicted sites can likely be increased further while maintaining high levels of accuracy. Interestingly, despite predicting a smaller number of m6A sites, many of the m6A predictions from xPore have not been reported by other protocols. Therefore direct RNA sequencing not only provides a simplified approach to profiling RNA modifications but also identifies novel sites that are possibly missed by complementary approaches.

Direct RNA-Seq has been used to analyse m6A in yeast (Garalde et al. 2018; H. Liu et al. 2019), Arabidopsis (Parker et al. 2020), RNA virus genomes (Kim et al. 2020) and in human cells (Workman et al. 2019; Leger et al. 2019; Lorenz et al. 2020). Here we showed that differential RNA modifications can be identified across a larger number of genetically diverse human cancer cell lines. Interestingly, even for m6A sites which were significantly different across cell lines, we found that they still preserved a certain level of m6A, suggesting that estimation of the modification rate is the key to understanding the dynamics of m6A. With direct RNA-Sequencing becoming widely available, we propose that the differential modification analysis can complement the differential expression analysis and provide insights into the highly complex landscape of RNA modifications and their roles in diseases.

## Methods

### xPore: A multi-sample two-Gaussian mixture model for identification of differential modifications

When a modified k-mer is passed through the pore, it causes the current intensity to differ from its canonical counterpart, enabling the detection of modification at the signal-level from dRNA-seq data. Based solely on current intensity levels, we aim to identify differentially modified positions between samples quantitatively.

Modelling a collection of transcripts at a single genomic site, we assume two distributions to correspond to the unmodified and modified RNAs that are shared across samples and allow the individual reads to fit both distributions with different degrees. With this assumption, we extend a standard two-Gaussian mixture model to support multiple sample comparison simultaneously. The multi-sample two-Gaussian mixture model allows the signal properties (i.e. means and variances) denoting the modified and unmodified RNAs to be shared across samples, yet accommodates sample-specific mixing weights, one of which is later used as an estimate of modification rate per sample.

Let *y*_*mn*_ be an intensity level mean of a read *n* from a sample *m* aligned at a given position *i* corresponding to a 5-mer *k*. We assume that the intensity level of a modified read is independently drawn from a normal distribution of modified *k* with a mean *μ*_*mod*_ and a variance 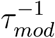, otherwise from another normal distribution of unmodified *K* with a mean *μ*_*unmod*_ and a variance 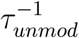. We denote *z*_*mn*_ to be 1 if a read is modified and 0 otherwise. Therefore, the conditional likelihood of all reads *y =* {*y*_1_,…, *y*_*mn*_} at a position *i* given *z =* {*z*_1_,…, *z*_*mn*_}, μ = {*μ*_*unmod*,_ *μ*_*mod*_}, *τ* = {*τ*_*unmod*,_ *τ*_*mod*_} can be written in the form of a mixture of the two Gaussians as follows,

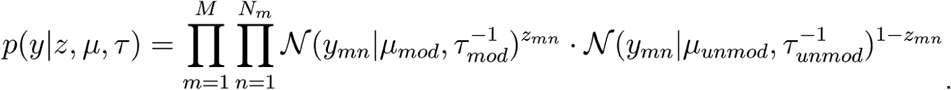

As a theoretical signal distribution of the unmodified k is available, we allow the model to favour the unmodified *k* unless the data reveal otherwise by imposing a Normal-Gamma distribution as a prior:

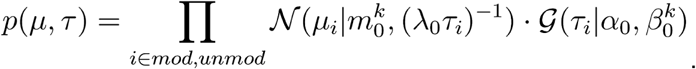

With this prior, the Gaussian parameters are regularised, which can help the inference for those positions that have a low number of reads.

We assume *z*_*mn*_ to follow a Bernoulli distribution with a probability *w*_*m*_, to which we refer as a modification rate of sample m:

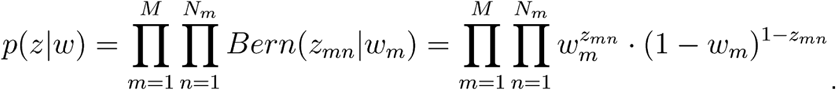

We also put a symmetric, uninformative Beta distribution as a prior on each sample modification rate as follows,

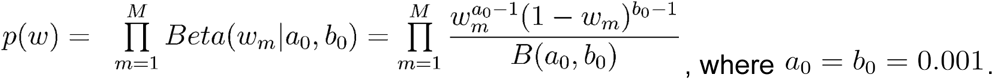

In order to make inference, we employ a variational Bayesian algorithm to estimate all model variables iteratively. The updating equations can be found in Supplementary.

### Post-modelling filter

At the positions where only a significant fraction of reads are modified, two Gaussians were well-distinguishable. On the other hand, at the sites where either complete modification or none is found, one of the distribution means is to converge into its prior with a large variance, forming an uninformative distribution. Reads from all conditions tested are modelled in the other distribution, indicating no differences among conditions. To discriminate those positions, we first tested the separation of the two distributions by calculating the probability of the overlapping area of the two clusters. We considered those positions with no more than 50% overlapping area as distinguishable. We also removed those positions where one is inside in the other, that is, when more than one intersection point has the density value higher than 0.1. We allow these thresholds to be adjustable by users.

### Statistical test for the differential modification rate (DMR)

Among the remaining sites, we assign the closer mean to the prior to be the unmodified and the other to the modified RNAs. For each condition, the model yields a modification rate for each sample. To prioritise positions with differentially modified in a pairwise comparison, we performed a two-tailed, unpooled z-test on the modification rate difference of any two conditions for each position:

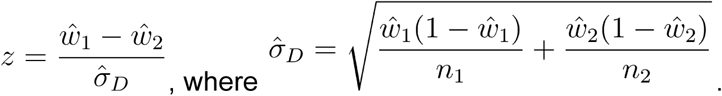

In case replicates are available, the average of the read coverages and the modification rates across the replicates for each condition are used to compute the *n*_1,_ *n*_2_ and *ŵ*_1_, *ŵ*_2_ respectively. As a result, the z-scores were used to rank the differential modification positions.

### Profiling m6A validation set

We created a validation case-control set of transcriptome-wide m6A modification at single-base-resolution by applying m6A-Crosslinking-Exonuclease-sequencing (m6ACE-seq) to quantify relative differences in methylation levels between wild-type and *Mettl3*-knockout HEK293T RNA samples (Koh, Goh, and Sho Goh 2019). We identified METTL3-dependent m6A sites using previously-determined criteria (Koh et al 2019). Briefly, these sites exhibited a wild-type/Mettl3-knockout relative methylation level ratio >=4.0 (p-value of one-tailed t-test < 0.05). As a result, a comprehensive profile of 15,703 genomic positions with significantly differential m6A modification was generated, covering 4,508 unique genes.

### Data generation and pre-processing

Following the standard steps, the ionic current readout for each FAST5 file was basecalled using Guppy and stored in FASTQ files (nf-core/nanoseq: http://doi.org/10.5281/zenodo.3697960).

Although both signal and sequence are generated from the sequencing machine and its proprietary basecaller software, segmenting the contagious signal *samples* into *events* and assigning each to a k-mer is an essential prerequisite for raw signal analysis. It is assumed that a single event comprises a set of samples drawn at the time when a k-mer of a strand resides in the pore, and the consecutive event when the strand moves past the pore at another base, where k is five for an RNA strand.

In achieving this, we applied “Nanopolish-Eventalign” to associate signal partials with their corresponding reference nucleotides (Loman, Quick, and Simpson 2015; Simpson et al. 2017). As a result, each read is segmented and its properties including reference k-mer, model k-mer events and their corresponding signal segments with the observed and expected normalised means along were reported.

However, the results from Nanopolish multiple events aligned to the same position, model k-mer was not matched to the reference k-mer, and were skipped. We handled these conditions by combining those events aligned to the same position in the strand and recalculating the average normalised 5-mer event means weighted by their event length. We ignored those skipped positions, discarded those mismatched k-mers. For sufficient coverage, moreover, only positions with 30-1,000 reads aligned were considered. In total, we modelled 9,509,290 genomic positions, involving 5,621 genes.

## Code availability

Our implementation in Python is available at https://github.com/GoekeLab/xpore.

